# A new method for obtaining bankable and expandable adult-like microglial cells

**DOI:** 10.1101/2021.03.31.437811

**Authors:** Min-Jung You, Chan Rim, Youn-Jung Kang, Min-Soo Kwon

**Affiliations:** Department of Pharmacology, Research Institute for Basic Medical Science, School of Medicine, CHA University, CHA BIO COMPLEX, 335 Pangyo, Bundang-gu, Seongnam-si, Gyeonggi-do, 13488, Republic of Korea; Department of Biochemistry, Research Institute for Basic Medical Science, School of Medicine, CHA University, CHA BIO COMPLEX, 335 Pangyo, Bundang-gu, Seongnam-si, Gyeonggi-do, 13488, Republic of Korea

**Keywords:** Microglia, Neuroepithelial layer, subculture, banking

## Abstract

The emerging role of microglia in neurological disorders requires a novel method for obtaining massive amounts of adult microglia because current *in vitro* methods for microglial study have many limitations, including a limited proliferative capacity, low cell yield, immature form, and too many experimental animals use. Here, we developed a new method for obtaining bankable and expandable adult-like microglial cells using the head neuroepithelial layer (NEL) of mouse E13.5. The NEL includes microglia progenitors that proliferate and ramify over time. Functional validation with a magnetic-activated cell sorting system using the NEL showed that the isolated CD11b-positive cells (NEL-MG) exhibited microglial functions, such as phagocytosis (microbeads, amyloid β, synaptosome), migration, and inflammatory changes following lipopolysaccharide (LPS) stimulation. NEL was subcultured and the NEL-MG exhibited a higher expression of microglia signature genes than the neonatal microglia, a widely used *in vitro* surrogate. Banking or long-term subculture of NEL did not affect NEL-MG characteristics. Transcriptome analysis revealed that NEL-MG exhibited better conservation of microglia signature genes with a closer fidelity to freshly isolated adult microglia than neonatal microglia. This new method effectively contributes to obtaining adult-like microglial cells, even when only a small number of experimental animals are available, leading to a broad application in neuroscience-associated fields.

## Introduction

Since their discovery in the 20th century by Rio-Hortega, microglia were considered professional phagocytes, similar to macrophages for a long time. However, recent studies propose microglia as a promising target beyond phagocytes under neuroinflammatory/neurodegenerative conditions (Nayak, Roth, & McGavern, 2014). Microglia are capable of morphological remodeling without any indication of insult or neurodegeneration (Nimmerjahn, Kirchhoff, & Helmchen, 2005), suggesting a broad functional repertoire, including the maintenance of biochemical homeostasis, neuronal circuit maturation during development, and experience-dependent remodeling of neuronal circuits in the adult brain (Lenz & Nelson, 2018). The dynamic microglial function lasts during our entire lifetime, and its disturbance can induce abnormal neurodevelopment and neurodegeneration (Matcovitch-Natan et al., 2016).

Despite the increase in knowledge regarding microglial biology (Prinz, Jung, & Priller, 2019), lack of a well-validated *in vitro* culture method remains a bottleneck for establishing a therapeutic strategy targeting microglia. The current *in vitro* culture of microglia still requires a microglial cell line and primary culture using neonatal cortex or the adult brain. Microglial cell lines, such as BV2, N9, SIM-A9, Mocha, and MHC3, have an advantage in proliferation and subculture; however, they differ from adult microglia in genetic and functional aspects due to immortalization (Timmerman, Burm, & Bajramovic, 2018). Neonatal microglia have not been artificially manipulated, but have disadvantages regarding their proliferative capacity, and subculture and banking abilities. Both the microglial cell line and neonatal microglia rarely express key adult microglia genes; in particular, TMEM119 immunoreactivity (IR) is missed (Bennett et al., 2016; Butovsky et al., 2014) because microglial cells develop according to a stepwise program (Matcovitch-Natan et al., 2016). Acute isolation of adult microglia is challenging due to their restricted proliferative capacity, cell viability, and requirement of a greater number of animals (Butovsky et al., 2012; Timmerman et al., 2018). The iPSC-derived microglia-like cells might also be an alternative method; however, this method is expensive, time-consuming, and not easily accessible (Muffat et al., 2016). In addition, iPSC-derived microglia-like cells do not originate in the yolk sac and do not show adequate TMEM119 IR compared to adult microglia (Abud et al., 2017). Thus, while the demand for microglia for use in research is increasing, there are no current *in vitro* methodologies that overcome the above-mentioned limitations.

Mouse microglia progenitors begin to migrate from the yolk sac into the CNS at E8.5 and surround the neuroepithelial layer (NEL) in the part corresponding to the head (Hoeffel et al., 2015; Nayak et al., 2014). Subsequently, microglia progenitors move into the CNS parenchyma until approximately E18.5 and are matured in the CNS microenvironment (Gosselin et al., 2017). As the CNS matures, microglia acquire a ramified morphology, surveying the surrounding parenchyma via movement of dynamic processes (Nimmerjahn et al., 2005).

In the support of the idea that microglia progenitor derived from yolk sac should pass NEL to enter CNS parenchyma, we could obtain and co-culture microglial progenitors and neuroepithelial cells together by dissecting mouse E13.5 head NEL, expecting that neuroepithelial cells play a role as feeder cells. By this new methodology, we could establish *in vitro* approach to obtain bankable and expandable adult-like microglial cells.

## Results

### Expandable and ramified microglial cells are generated from mouse E13.5 head neuroepithelial layer

To determine the optimal period for obtaining a high yield of microglia progenitors in the head NEL, we dissected head NEL from mouse E9.5, E13.5, and E17.5 and cultured them (Fig. 1A). Immunofluorescence analyses revealed that the highest numbers of microglia progenitors were obtained from the mouse E13.5 NEL compared to that of mouse E9.5 or E17.5 (Fig. S1). The flow cytometry results showed that E13.5 NEL contain more CD11b-positive cells than the brain cortex at E13.5 (Fig. S1). We next examined whether microglia progenitors were expandable and ramified over time in this culture system. The proportion of CD11b-positive cells increased from 14.2% up to 54.0% after 21 days of culture (Fig. 1B). The immunofluorescence results showed that the number of microglial cells that were IBA-1-, PU1-, and F4/80-positive increased over time. The percentage of double positive cells (Ki67 and IBA-1) also increased to 50%, suggesting that microglia have a high proliferative capacity in this culture system (Fig. 1C).

**Figure 1.**
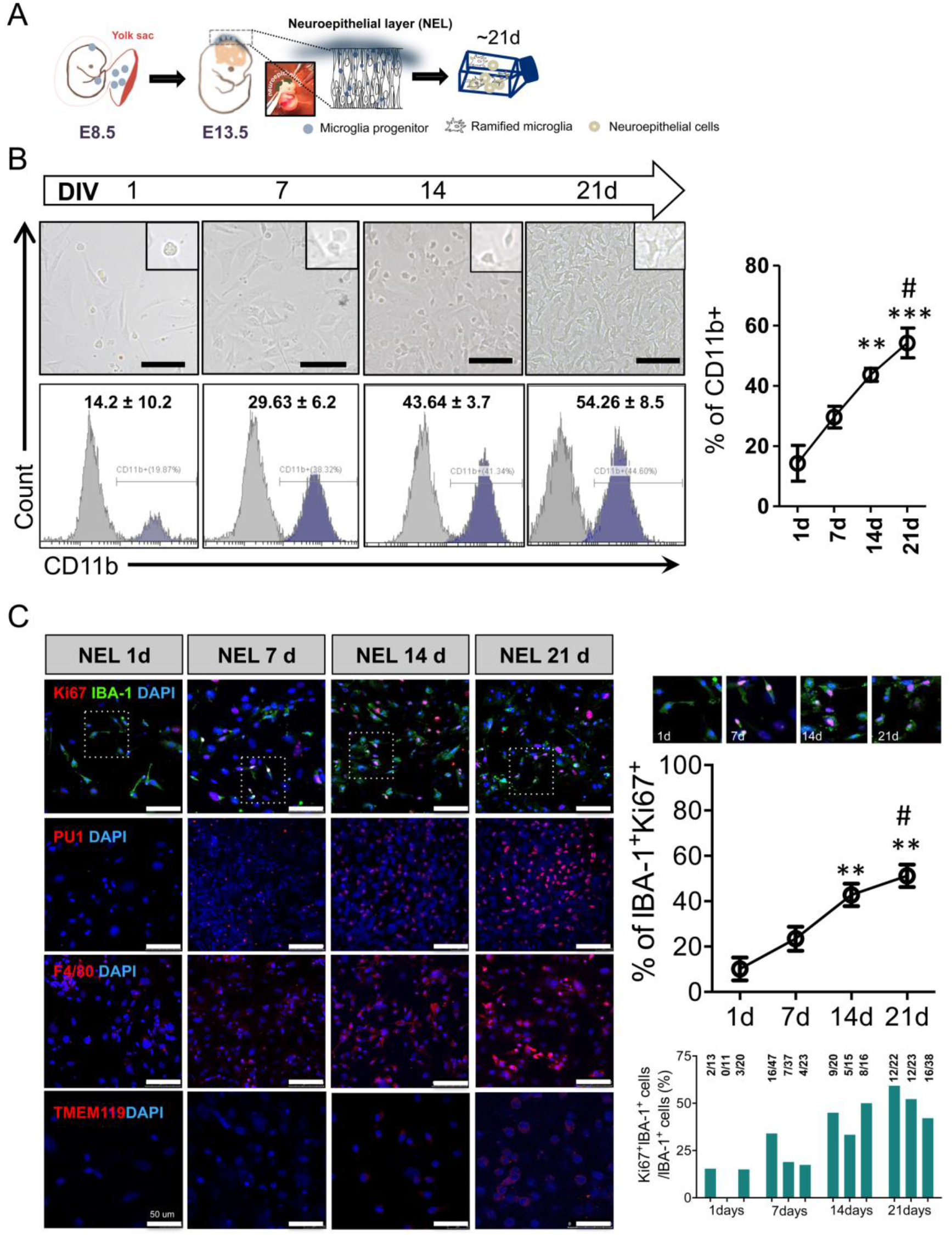
Expandable microglia generation from the embryo neuroepithelial layer. (A) Mouse head neuroepithelial layer (NEL) was dissected at E13.5 and was cultured for 21 days (d). (B) Microglia progenitors proliferated and ramified over time. (C) IBA-1^+^Ki67^+^ cells were stained and the ratio of IBA-1^+^Ki67^+^ cells/IBA-1^+^ cells increased over time. The number of PU1- and F4/80-positive cells also increased with the progression of the culture. TMEM119 was stained weakly at 21 d. Scale bar = 100 μm. Proliferative microglia (IBA-1^+^Ki67^+^ cells) were counted using microscopy. Sampled areas were selected randomly from 100× fields from three independent experiments. A *post-hoc* test was conducted using Tukey’s multiple comparison tests. ^**^p < 0.01, ^***^p < 0.001 compared to 1 d, ^#^*p* < 0.05 compared to 7 d.

### Functional validation of magnetically isolated CD11b positive cells (NEL-MG)

CD11b-positive cells were isolated using a magnetic-activated cell sorting (MACS) system from the E13.5 NEL (NEL-MG). NEL-MG stained positive for microglia markers with IBA-1 and CX3CR1. NEL-MG showed a relatively weak TMEM119 IR; however, they did not express Ki67 (Fig. 2A). Next, we conducted microglia-specific functional assays, including phagocytosis, migration, and cytokine/chemokine release using NEL-MG. NEL-MG showed phagocytic function against stimuli, such as synaptosome, amyloid β, and microbeads (Fig. 2B). The scratch wound assay results showed that NEL-MG could migrate (Fig. 2C). Based on the cytokine and chemokine profiles, unstimulated NEL-MG released detectable cytokines and chemokines, including the C-X-C motif chemokine ligand 10 (CXCL10), CXCL1, C-C motif chemokine ligand 12 (CCL12), CCL3, TNF-α, and IL-6. The addition of endotoxin (LPS) triggered the release of chemokines and cytokines above baseline levels (untreated) (Fig. 2D), which was verified at the transcriptional level, with an increase in the mRNA expression of iNOS, TNF-α, IL-1β, IL-6, and CCL3 (Fig. 2E). Therefore, NEL-MG acted as a substrate for studying the functional changes in microglia to screen for inflammatory modulators.

**Figure 2.**
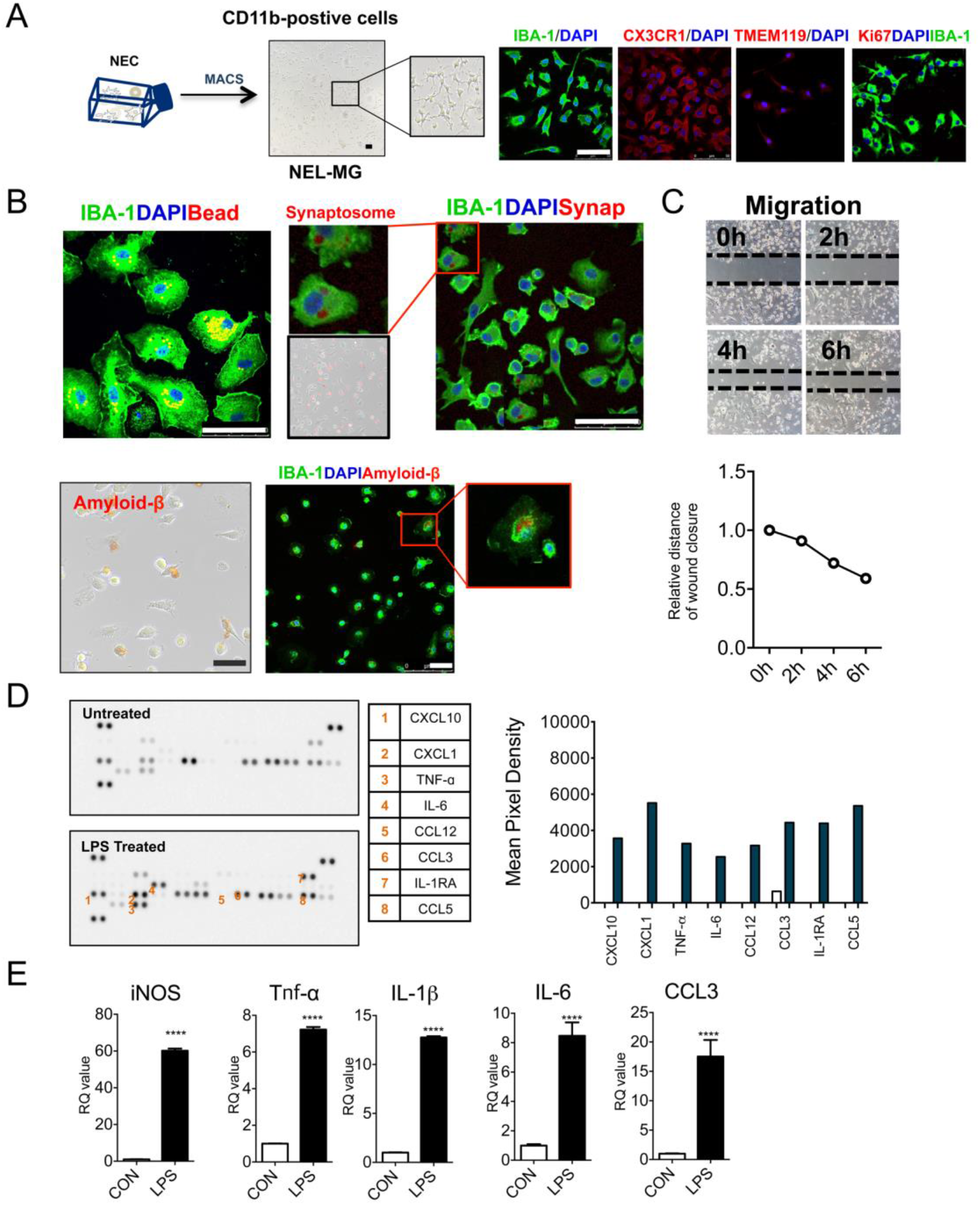
Functional validation of NEL-MG. (A) CD11b^+^ cells were isolated from cultured NEL for 21 days using an MACS system (NEL-MG) and NEL-MG showed IBA-1, CX3CR1, and TMEM119 IR, but not Ki67 IR. (B) A phagocytosis assay using microbeads, synaptosome, and amyloid β was used to assess NEL-MG. (C) Migration performance was assessed using a wound healing assay. (D) Cytokines and chemokines released in the supernatant of NEL-MG in response to LPS challenge. (E) Transcription in NEL-MG as a baseline (CON) and after LPS stimulation (LPS) (n = 3). The experiment was conducted independently three times. Scale bar = 50 μm. An unpaired t-test was conducted to compare the two groups. ^****^p < 0.0001 compared to the control.

### Long-term passage culture of NEL and a higher expression of NEL-MG on microglia signature genes

After validating the microglial function of NEL-MG, we hypothesized whether the long-term passage culture of E13.5 NEL was possible. Whenever the cells reached stratification, the cells in one 25T flask were subcultured into two 25T flasks and cultured for approximately 10 days. As we undertook this passage culture, we obtained approximately double the microglial cells until at least passages 6–7 (first obtained microglial cell number × 2^n^, n = passage number). This was because we could obtain about 50% of CD11b-positive cells from NEL, despite the long-term passage culture (Figs. 3A and S2). However, the required time for the next passage gradually increased as the subculture progressed and subculture without obtaining twice as many cells only occurred after passage 7. Therefore, the total number of microglia obtained when we used a cutoff of 100 days was 30 times higher than using the neonatal microglia method (Fig. S2).

**Figure 3.**
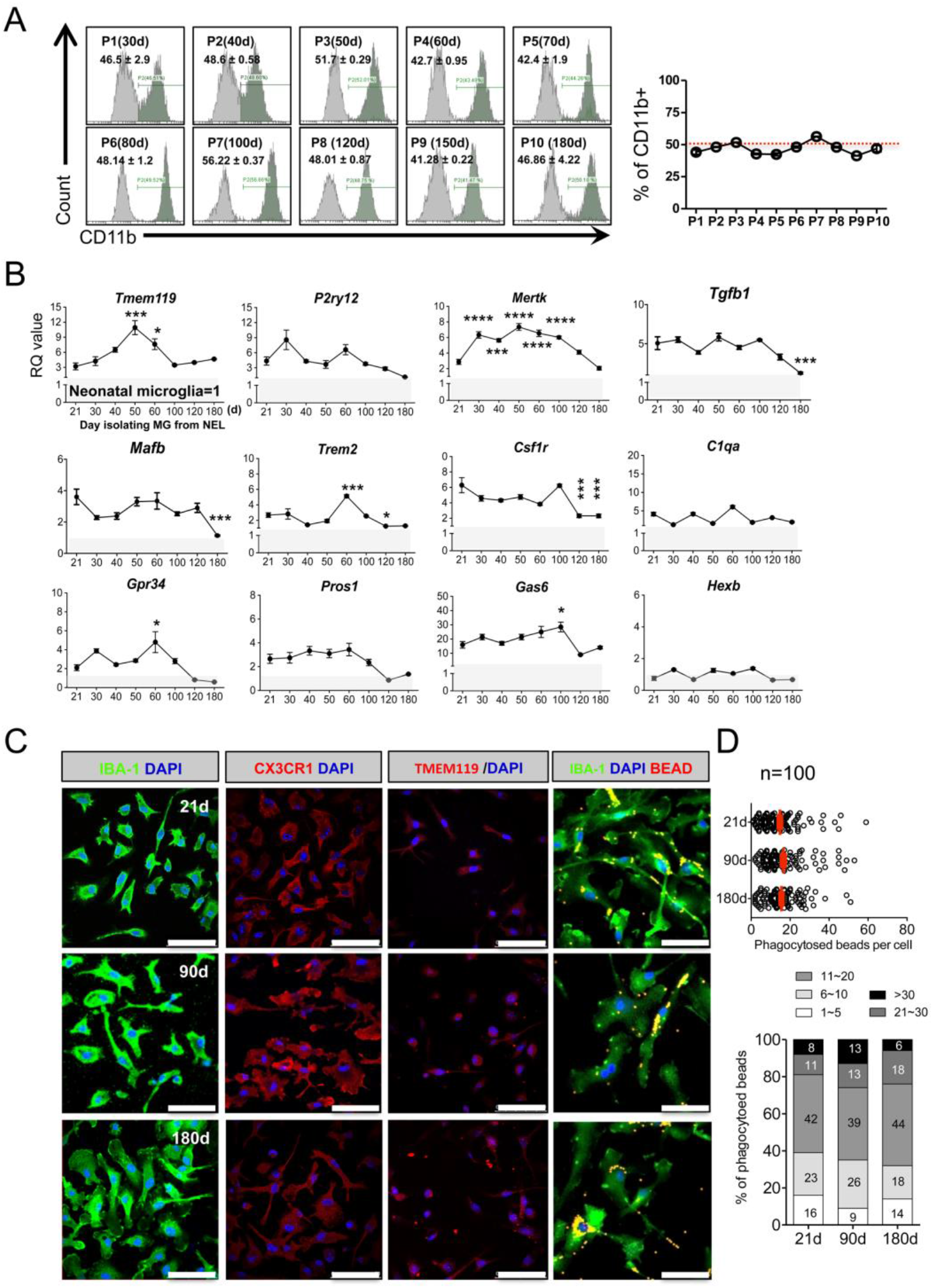
Stable maintenance and superiority of NEL-MG on adult microglial genes despite the long-term passage culture. When NEL reached stratification, cells in one 25T Flask were divided into two 25T flasks, and then cultured for 21 days (d) for flow cytometry. (A) The ratio of CD11b^+^ microglia among NEL was approximately 50%, showing a stable maintenance ratio, despite the long-term passage culture (n=3, mean ± SD) in flow cytometry. (B) NEL-MG were isolated based on the NEL culture time (21, 30, 40, 50, 60, 100, 120, and 180 d) and alterations in the adult microglial genes were examined compared to neonatal microglial cells. The experiment was performed three times independently (n = 3 per group) and the data represents mean ± standard error of the mean (SEM). A *post-hoc* test was conducted using Dunnett’s multiple comparison test. ^*^p < 0.05, ^***^p < 0.0001, ^****^p < 0.00001 compared to the 21-d group. Gray zone represents a value = 1 (neonatal microglia). (C) IBA-1, CX3CR1, and TMEM119 IRs and phagocytic function were not different in NEL-MG isolated from NEL at 21, 90, and 180 d. Scale bar = 50 μm.

We next examined the alterations in the microglial signature genes in NEL-MG obtained from long-term cultured NEL to determine the optimal period for obtaining adult-like microglia. NEL-MG were isolated from 21, 30, 40, 50, 60, 100, 120, and 180 days of NEL culturing, and the expression of microglial signature genes in each NEL-MG group was compared to that in neonatal microglia. The expression of most of the microglia identity genes was stably maintained and was higher than in neonatal microglia up to 180 days; however, the expression levels of *Tgfb1, Mafb, Trem2*, and *Csf1r* were significantly lower at 120 days than at 21 days (Fig. 3B). The expression level of the representative adult microglia gene, *Tmem119*, gradually increased until isolation from the 50-day cultured NEL and decreased afterward; however, it remained approximately 3-fold higher than *Tmem119* expression in neonatal microglia. Our data strongly suggest that adult microglia-like NEL-MG were obtainable from cultured NEL for 21–100 days (P0–P7). However, the IR of microglial markers, such as IBA-1, CX3CR1, and TMEM119, and their phagocytic function did not change, regardless of the NEL culture duration (Fig. 3C and D).

### Banked NEL-MG stably maintain the expression of overall microglia signature genes

Animals must be sacrificed to obtain primary microglia; therefore, whether NEL is bankable needs to be determined. NELs (1 × 10^7^ cells/mL) were banked for approximately ten months in media composed of Dulbecco’s modified Eagle medium (DMEM) with 20% fetal bovine serum (FBS) and 10% DMSO (banked NEL, bNEL). As shown in Figure 4A, after thawing, the bNEL proliferated well, similar to fresh NEL. To investigate whether the banking timing affected NEL-MG characteristics, NELs were banked at different passages (P1, P2, P3, and P4) and were cultured for 70 days after thawing. Each NEL-MG group isolated from bNEL at different time points (7, 14, 21, 40, and 70 days) showed a stable expression of microglia signature genes. The expression levels of these genes remained higher than those in neonatal microglia (Fig. 4B); however, *TMEM119* mRNA expression showed a decreasing trend at 70 days (not significant). *Csf1r* expression was also reduced after thawing; however, its mRNA level was gradually restored as the culture progressed. NEL-MG isolated from bNEL did not show alterations in the phagocytic function compared to NEL-MG isolated from fresh NEL (Figs. 3D and 4C).

**Figure 4.**
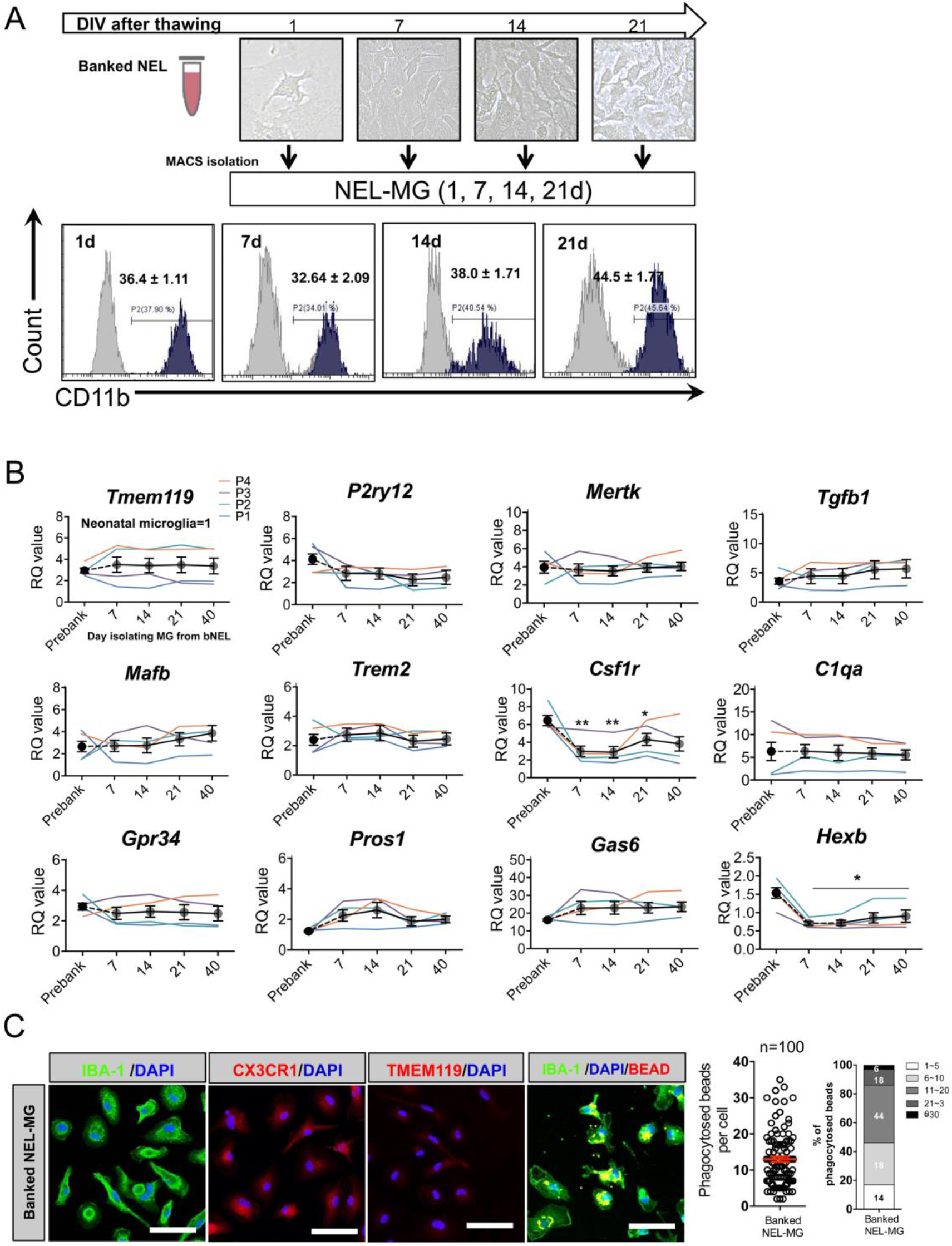
Validation of NEL-MG isolated from banked NEL. (A) NEL were banked for about 10 months and thawed. The thawed NEL were cultured for 1, 7, 14, 21, 70 days (d) and NEL-MG were isolated at each time point for flow cytometry. (B) Banked NEL at different passages (P1, P2, P3, and P4) were thawed and cultured for 7, 14, 21, 40, and 70 d. NEL-MG were isolated at each time point for qPCR (n = 4 per group) (black circle: mean value of P1, P2, P3, and P4) and the data represent mean ± standard error of the mean (SEM), ^*^p < 0.05, ^**^p < 0.001 compared to the pre-banked group. A *post-hoc* test was conducted using Dunnett’s multiple comparison tests. (C) Immunofluorescence study of IBA-1, CX3CR1, and TMEM119 IR and a phagocytosis assay were conducted using banked NEL-MG (70 d). Scale bar = 50 μm.

### Transcriptome analysis revealed that NEL-MG are closer to adult microglia than neonatal microglia

To more thoroughly characterize NEL-MG compared to the BV2 cell line, neonatal microglia, and adult microglia, we conducted transcriptome analysis. Using 3D principal component analysis (PCA) of the whole transcriptome, each group was classified into distinct clusters based on their identity. NEL-MG and bNEL-MG were clustered together and were distinct from the BV2 cell line and neonatal microglia. The component explaining the majority of the dataset variance was PC1, which most prominently distinguished neonatal microglia and NEL-MG from adult microglia and occupied an edge position along PC1. The second principal component (PC2) uniquely distinguished the BV2 cell line from neonatal microglia and adult microglia. The third principal component (PC3) uniquely distinguished NEL-MG from neonatal microglia (Fig. 5A). The PCA Euclidean distances to the adult microglia were calculated between all pairs of points in each group, using PC1, PC2, and PC3 on the microglia signature genes (Table S1) (Butovsky et al., 2014; Najafi et al., 2018). NEL-MG and bNEL-MG were significantly closer to adult microglia than BV2 and neonatal microglia (Fig. 5B). The hierarchy analysis of the microglia signature genes showed that NEL-MG and bNEL-MG formed a cluster and had more similarities to adult microglia than to BV2 cell line and neonatal microglia (Fig. 5C). The overall expression changes (z-score normalized) in microglia signature genes were compared among the BV2 cell line, neonatal microglia, NEL-MG, and adult microglia (mean ± standard deviation [SD]). Both NEL-MG and adult microglia were localized in the opposite area (positive Z-score) from neonatal microglia and the BV2 cell line (negative Z-score). The overall expression of microglia signature genes in the NEL-MG was elevated to the same expression level as that of the genes in adult microglia (Fig. 5D). In a normalized gene expression plot, the scatters in NEL-MG were more overlapped and closer to the diagonal line than neonatal microglia (Fig. S3).

**Figure 5.**
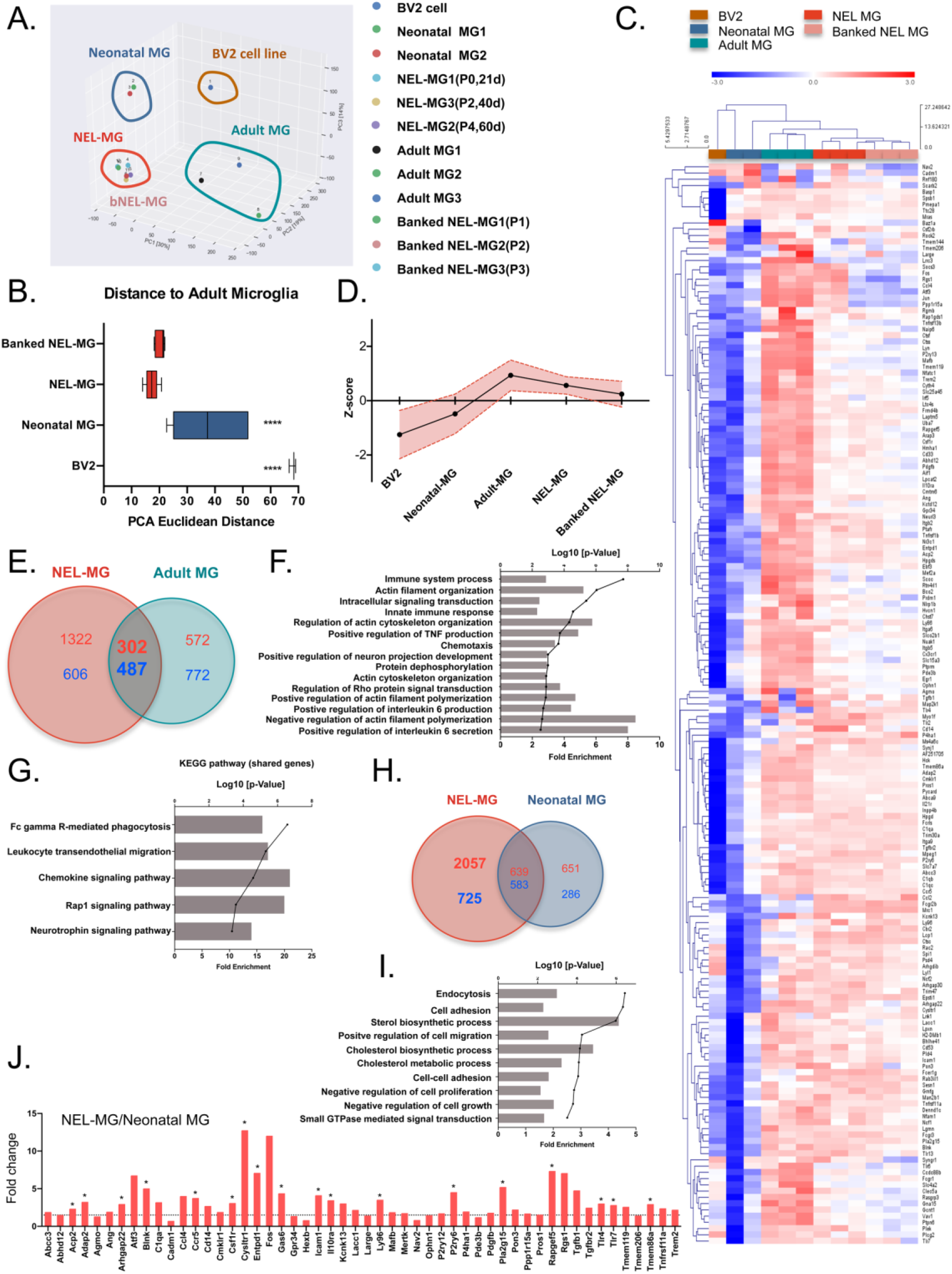
NEL-MG with a closer fidelity to adult microglia. (A) Principal component analysis (PCA) was performed to determine the similarity among the BV2 cell line, neonatal microglia (MG), NEL-MG, banked NEL-MG (bNEL-MG), and adult MG. (B) PCA Euclidean distances to the adult MG were calculated between all pairs of points in each group using PC1, PC2, and PC3 on microglial signature genes. ^****^p < 0.00001. (C) Clustering heatmap representation of microglia signature gene expression between clusters. The scale represents the median absolute deviation by row. For calculating the distance, a Euclidean distance metric and average linkage clustering were performed using MeV software. (D) The overall expression changes (z-score normalized) were plotted among groups as mean ± SD. (E) Differentially expressed genes between NEL-MG and adult MG. The number of shared genes was 789. (F) Gene ontology (GO) analysis of the shared genes (789 genes). (G) KEGG pathway analysis of the shared genes (789 genes). (H) Differentially expressed genes between NEL-MG and neonatal MG. The number of unshared genes in NEL-MG was 2,782 (I). GO analysis of the 2,782 genes. (G) KEGG pathway analysis of the 2,782 genes. (J) Upregulated differentially expressed genes between NEL-MG and neonatal MG relative to adult MG were compared regarding the microglial signature genes. The dotted line indicates 1.5-fold change compared to neonatal MG. ^*^ p < 0.05.

We next performed differential gene expression analysis (DEG, absolute fold changes > 1.5, p < 0.05) and identified that NEL-MG shared 789 genes (up DEG: 302, down DEG: 487) with adult microglia (Fig. 5E). Gene ontology (GO) analysis revealed the gene subsets of the immune system process, actin filament organization, intracellular signaling transduction, and innate immune response (Fig. 5F). The KEGG pathway of the shared genes indicated Fc gamma R-mediated phagocytosis, leukocyte transendothelial migration, and chemokine signaling pathway (Fig. 5G). NEL-MG showed 2782 DEG (2,057 upregulated and 725 downregulated) compared to neonatal microglia (Fig. 5H), and the GO analysis revealed gene subsets, including those involved in endocytosis, cell adhesion, cell migration, cholesterol biosynthetic process, cholesterol metabolic process, and the negative regulation of cell proliferation (Fig. 5I). To better understand the differential expression level of microglia signature genes between NEL-MG and neonatal microglia, we compared the fold change of the expression of microglia signature genes (Table S1), and 18 genes were significantly changed (Fig. 5J).

### Comparison of NEL-MG with adult microglia regarding microglia signature genes

To further confirm the transcriptome analysis, we compared the mRNA expression of microglial signature genes among the BV2 cell line, SIM-A9 cell line, neonatal microglia, NEL-MG (50 days), and acutely isolated adult microglia. qRT-PCR analyses demonstrated that NEL-MG had higher or similar expression levels of microglia signature genes, including *Tgfb1, Tgfbr1, Trem2, Csf1r, C1qa, Pros1*, and *Gas6* (Fig. 6A). Although *Tmem119* and *P2ry12* expression levels of NEL-MG did not reach those in adult microglia, they were higher than in neonatal microglia. In addition, the intensity of TMEM119 IR in NEL-MG (50 days) was similar to that in adult microglia, whereas BV2 and neonatal microglia did not show TMEM119 IR (Fig. 6B).

**Figure 6.**
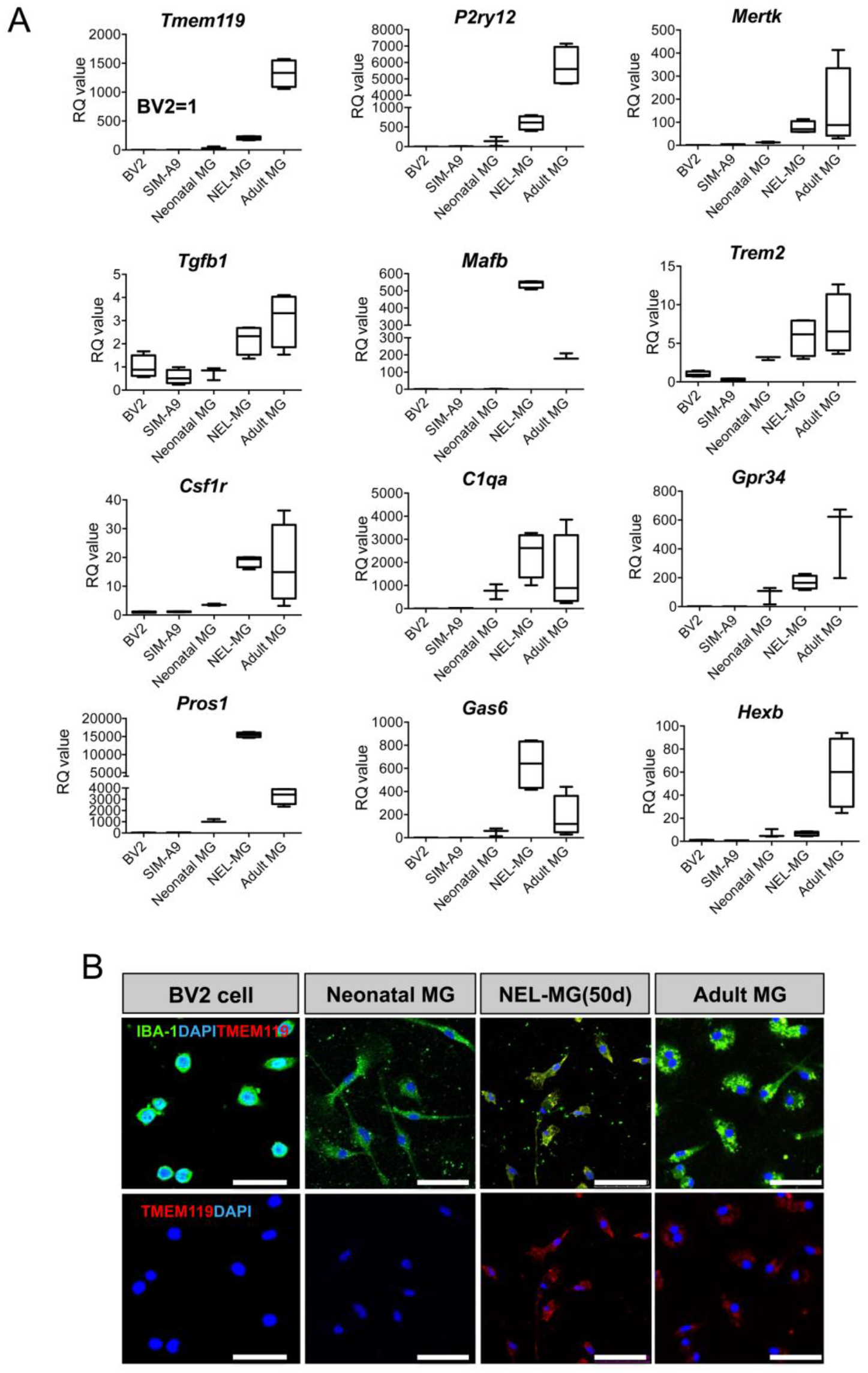
Validation of transcriptome analysis. (A) The expression level of adult microglial genes was compared among the BV2 cell line, SIM-A9, neonatal microglia (MG), NEL-MG (50 d), and adult MG. n = 3–4 as independent experiments. (B) IBA-1 and TMEM119 were stained in NEL-MG and adult MG, but not in neonatal MG. Scale bar = 50 μm

### Cultured NEL-MG showed lower re-expansion on neuroepithelial cells

To further explore whether cultured NEL-MG could be proliferative again on neuroepithelial cells as a feeder, we isolated and cultured GFP-expressing NEL-MG derived from NEL of GFP mice (GFP-expressing NEL-MG). Then, GFP-expressing NEL-MG were cultured on the remaining neuroepithelial cells (from B6 mice) after NEL-MG isolation. As shown in Figure 7A and B, using flow cytometry, we confirmed that the relative ratio of GFP-expressing NEL-MG did not increase significantly over time, although GFP^+^Ki67^+^ cells were examined in the immunofluorescence study. Therefore, NEL-MG had a lower proliferative capacity, even when they were plated again on neuroepithelial cells.

**Figure 7.**
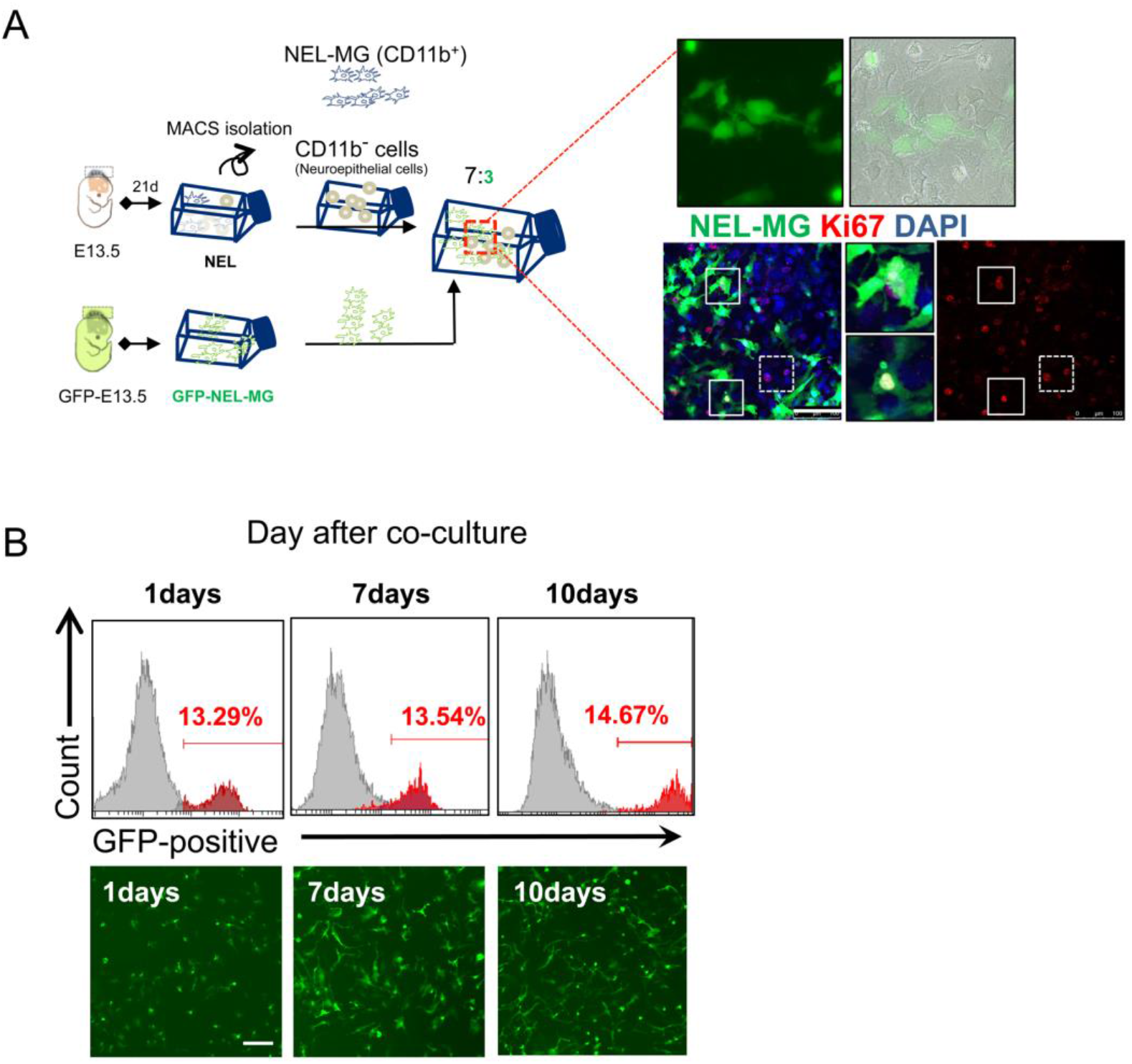
NEL-MG showed a low re-expansion capacity. (A) GFP-expressing NEL-MG were plated on a neuroepithelial cell feeder and cultured. GFP-expressing NEL-MG were obtained from B6-EGFP mice using our methodology. GFP-expressing Ki67-positive NEL-MG (white line square) or no GFP-expressing Ki67-positive cells (neuroepithelial cells, dot line square) were examined in the immunofluorescence study. (B) According to the flow cytometry results, the number of GFP-positive cells was not changed over the culture duration, but the morphological changes to the ramified form were examined over time. Scale bar = 100 μm.

## Discussion

Our robust protocol allowed us to obtain bankable and expandable microglial cells, leading to the production of a high yield of microglial cells that were closer to adult microglia than neonatal microglia based on PCA. We based our study on the developmental process, in which mouse microglial cells were derived from yolk sac macrophage precursors and colonized into nascent parenchyma via the head neuroepithelial layer between E8.5 and E18.5 (Nayak et al., 2014). The neuroepithelial layer in this area includes mainly two types of cells: neuroepithelial cells and microglia progenitors. Thus, we obtained microglial cells by dissecting and culturing the neuroepithelial layer cells with lower cell heterogeneity than the brain cortex, which is composed of neurons, microglia, astrocytes, and oligodendrocytes, making it possible to obtain purer microglial cells. Neuroepithelial cells were assumed to act as feeders for microglia proliferation and maturation, which was evidenced by our data, showing that microglia progenitors proliferate with Ki67-positive labeling on neuroepithelial cells, although NEL-MG alone failed to proliferate. Therefore, neuroepithelial cells within a co-culture system play a critical role as feeders for microglial proliferation and maturation. This result also indicates that the matured form of NEL-MG does not have the same proliferative capacity as microglia progenitors, despite the neuroepithelial support. Similarly, an astrocyte feeder layer or an astrocyte-conditioned medium makes microglia morphology and other properties resemble adult microglia more closely (Bohlen et al., 2017; Tanaka &Maeda, 1996). Compared to the astrocyte feeder method, our proposed protocol has advantages in a simpler methodology making banking and subculture possible.

Microglia rely on sustained CSF-1 stimuli, which is a well-established trophic cue for survival (Elmore et al., 2014), and TGF-β family members promote microglial survival and development (Butovsky et al., 2014). Along with CSF-1 and TGF-β, cholesterol is another main factor that permits the robust survival of highly ramified adult microglia (Bohlen et al., 2017). Our GO results indicate that NEL-MG shared pathways in general microglia function with adult microglia compared to neonatal microglia. However, NEL-MG was distinct from neonatal microglia, especially regarding the cholesterol biosynthetic process and metabolic process. Given the link between an inadequate delivery of cholesterol and generalized microglial dysfunction, the cholesterol metabolic system might be involved in key features in a more matured form of NEL-MG than neonatal microglia. The expression of microglial signature genes, including *TMEM119, P2RY12*, and *CX3CR1*, is gradually reduced with age or in neurodegenerative diseases, suggesting the loss of homeostatic microglial function (Deczkowska, Amit, & Schwartz, 2018). *De novo* synthesized cholesterol transported via APOE- or APOJ-containing nano-discs within the CNS (Pfrieger & Ungerer, 2011; Rapp, Gmeiner, & Huttinger, 2006) contribute to amyloid β clearance and microglial cholesterol content (Lee, Tse, Smith, & Landreth, 2012). Increased lipid metabolism is required to fuel protective cellular functions, such as phagocytosis (Loving & Bruce, 2020). Thus, we speculate that alterations in the cholesterol metabolic system might be associated with the characteristics of the matured form of NEL-MG, which is closer to adult microglia.

There are several aspects that require further exploration regarding NEL-MG. First, the cultured NEL-MG had a lower proliferative capacity despite the re-support of neuroepithelial cells. Microglial cells are dedifferentiated when they lose their cell– cell interactions. Acutely isolated adult microglial cells rapidly lost their identity in culture media, suggesting microglial dedifferentiation (Bohlen et al., 2017). NEL-MG might also be dedifferentiated when they are separate from neuroepithelial cells, which might be associated with a reduced proliferative capacity. Second, the supportive capacity of neuroepithelial cells as a feeder gradually decreased as the passage culture of NEL progressed, leading to a decrease in the microglial proliferative capacity. Third, the expression of *hexosaminidase subunit beta* (*Hexb*), which is a microglial core gene stably expressed despite the pathological status (Masuda et al., 2020), and *colony stimulating factor receptor 1* (*Csf1r*) was reduced in bNEL-MG compared to pre-banked NEL-MG and there was no difference in *Hexb* expression between NEL-MG and neonatal microglia. To overcome these limitations, further studies are required to develop an advanced methodology to produce improved NEL-MG with defined factors, which can induce sustained proliferation, maturation, subculture, and banking, instead of using neuroepithelial cell feeder.

Regarding the applications of our method, we propose the following as a platform for future use in the neuroscience field: 1) mass production of microglial cells for high-throughput drug screening, and 2) rapid and efficient generation of mutant microglial cells. Compared to neonatal microglia culture, our proposed method enabled us to obtain approximately 30 times more microglial cells when we used a cutoff of 100 culture days. Drug screening for inflammatory modulators targeting neuroinflammation requires significant amounts of microglial cells; however, the currently used *in vitro* method for primary microglia can barely meet the requirements due to limited cell yields and the number of animals required. Thus, our method contributes to overcoming the main limitation of the currently used technique. In addition, the introduction of a newly reported mutant gene into the NEL allowed the production of mass mutant NEL-MG. Using this methodology, we can rapidly evaluate the functional characteristics of mutant microglia without using a mutant generation. Given that the *in vivo* brain exhibits complex cell–cell interactions supported by contact-dependent signaling from the surrounding cells, mutant NEL-MG might be mixed with a brain organoid to further observe their behavior in a 3D brain environment, based on the fact that NEL-MG is mixed evenly with neurospheroids (Fig. S4) (Abud et al., 2017). In addition, NEL-MG might contribute to the development of a brain organoid with a controllable microglia ratio because the currently used brain organoid does not contain microglial cells, although one study has reported that a brain organoid might innately contain microglial cells (Ormel et al., 2018). Overall, our proposed method contributes to 3D brain culture and conventional 2D primary microglial culture.

In conclusion, we expect that our methodology contribute to overcoming the limitations of previous *in vitro* culture methods for microglia study by leveraging the NEL-MG platform, and increasing the adult-like microglial cell yield. Above all, we believe that our new methodology will reduce dramatically the use of experimental animals and increase the accessibility of microglial research.

## Material and methods

### Mouse embryonic neuroepithelial layer dissection and culture

Uteri from pregnant mice (female C57BL/6, Orient Bio Inc. Seoul, Korea or female B6-EGFP mice, Jackson lab) were dissected and soaked in Hanks’ Balanced Salt Solution (HBSS, Invitrogen, USA). The umbilical cord and yolk sac in the mesometrial surface of the uterus were removed using microsurgical instruments under a microscope and the embryo was taken out of the uterus. Then, the NEL was dissected carefully using a pair of microsurgical scissors. The shredded tissue was incubated with 1 mL of 1X Trypsin-EDTA (ThermoFisher, USA) for 3 min at 37 °C in a water bath. After centrifuging at 300 × *g* for 5 min, the cells were plated onto 25 cm^2^ flasks coated with poly-D-lysine (Sigma, MO, USA). The cells were cultured in DMEM-LG containing 10% FBS, 1X penicillin:streptomycin and 0.1% GlutaMAX at 37 °C in a 5% CO2 incubator. The culture medium was replaced with 5 mL of fresh growth medium after 24 h. Subsequently, half the culture medium volume was replaced with an equal volume of fresh growth medium twice per week.

### Isolation of microglial cells from NEL

On ∼day 21, when stratification was reached, NEL culture flasks were incubated with 1X trypsin-EDTA for 1 min, resulting in the detachment of an intact layer of cells in one piece. The pellet was resuspended in cold MACS buffer (containing a 1-volume dilution of PBS, 2 mM EDTA, and 0.5% BSA, pH 7.2), and myelin removal beads (Myelin Removal Beads II, 130-96-733, Miltenyi Biotec) were used according to the manufacturer’s protocol to prepare the cells for incubation with microbead-coupled anti-CD11b mAb. Briefly, cells were incubated with the beads at 4 °C for 15 min, and then the cells were washed onto the MS column on a magnetic separator. The column was washed thrice with PBS buffer, and the NEL-MG were obtained via positive selection. NEL-MG were resuspended in microglial complete culture medium (DMEM, 10% FBS, 0.1% GlutaMAX, 5 μg/mL insulin, and 1% penicillin/streptomycin), transferred to PDL-coated plates at a density of 2.5 × 10^5^ cells/mL, and cultured for subsequent molecular studies.

### Adult microglial cell isolation

Adult microglial cells were isolated from 8-10 week old mice (male C57BL/6, Orient Bio Inc. Seoul, Korea). Enzymatic cell dissociation was performed using an Adult Brain Dissociation Kit (130-107-677, Miltenyi Biotec) according to the manufacturer’s instructions. Brain tissue pieces (up to 500 mg) were transferred into a gentleMACS C tube (130-096-334, Miltenyi Biotec) containing 1,950 μL of enzyme mix 1 (enzyme P and buffer Z), and then 30 μL of enzyme mix 2 (enzyme A and buffer Y) was added. The gentleMACS C tube was tightly closed and attached upside down onto the sleeve of the gentleMACS Octo Dissociator with Heaters (130-096-427, Miltenyi Biotec), and the appropriate gentleMACS program (37C_ABDK_01) was run. After brief centrifugation to collect the samples at the bottom of the tube, the samples were filtered through a 70-μm strainer (130-098-462, Miltenyi Biotec), washed with D-PBS, and centrifuged again. The pellet was resuspended in cold D-PBS. The myelin and cell debris were removed using debris removal solution, followed by subsequent removal of erythrocytes using a red blood cell removal solution. Pure adult microglial cells were magnetically isolated using microbeads-coupled anti-CD11b mAb, as stated previously.

### Neonatal microglia culture

Postnatal 1–2-day-old B6 mice (Orient Bio Inc. Seoul, Korea) were sacrificed and soaked in 75% ethanol for 1 min. The cerebral hemispheres were dissected following standard techniques and anatomical landmarks, and the meninges were peeled off. The hippocampus, basal ganglion, and olfactory bulb were carefully removed using microsurgical instruments under a microscope, and the remaining cortical tissue was minced using a pair of microsurgical scissors. The shredded tissue was then incubated with 3 mL of HBSS (Invitrogen) for 5 min at 37 °C in a water bath. After centrifuging at 300 × *g* for 5 min, the cells were plated onto 75 cm^2^ flasks coated with poly-L-lysine (Sigma). Mixed glial cells were cultured in DMEM-LG containing 10% FBS and 0.1% GlutaMAX at 37 °C and 5% CO2 in an incubator. The culture medium was replaced with 15 mL of fresh growth medium after 24 h. Subsequently, half of the culture medium volume was replaced with an equal volume of fresh growth medium twice a week. Stratification was reached at the end of this period, and the microglial cells in the upper layer were harvested. On day 14, flasks were incubated with 1X trypsin-EDTA for 1 min, resulting in the detachment of an intact layer of cells in one piece. The pellet was resuspended in cold MACS buffer (containing 1-volume dilution of PBS, 2 mM EDTA, and 0.5% BSA, pH 7.2), and then myelin removal beads (Myelin Removal Beads II, 130-96-733, Miltenyi Biotec) were used according to the manufacturer’s protocol to prepare the cells for incubation with microbeads-coupled anti CD11b mAb. Briefly, the cells were incubated with the beads at 4 °C for 15 min, and then the cells were washed onto the MS column on the magnetic separator. The column was washed thrice with PBS buffer, and the magnetically labeled cells were obtained via positive selection. The cells were resuspended in microglial complete culture medium (DMEM, 10% FBS, 0.1% GlutaMAX, 5 μg/mL insulin, and 1% penicillin/streptomycin) and transferred to PDL-coated plates at a density of 2.5 × 10^5^ cells/mL for molecular studies.

### BV2 cell line and SIM-A9 culture

Murine BV-2 microglial cells were maintained in DMEM supplemented with 10% FBS and antibiotics at 37° C in a 5% CO2 incubator. Then, the cells were seeded onto 6-well plates at a density of 2 × 10^5^ cells/well for quantitative polymerase chain reaction (qPCR) analysis and RNAseq. SIM-A9 is a microglial cell line that was purchased from Kerafast (Boston, USA). These cells, referred to as SIM-A9 cells and related to native primary microglial cells, have been characterized for morphology and the release of cytokines/chemokines. After receiving, the cells were passaged in an uncoated 100 mm cell culture dish in DMEM/F-12 (Gibco, cat. # 11320-033) containing 10% heat-inactivated FBS (Gibco, cat. # 16000-044), 5% heat-inactivated horse serum (Invitrogen cat. # 16050-122), and 1% penicillin/streptomycin (Gibco, cat. # 15140122). Cells were cultured at 37 °C in an incubator with 5% CO2.

### Flow cytometry

The cells were collected and labeled using fluorochrome-conjugated monoclonal antibodies recognizing antigens (CD11b-PE, BD Biosciences #553311) at 4 °C for 15 min. After labeling, the cells were washed twice in PBS and resuspended in a final volume of 400 μL. Flow cytometry was performed using a CytoFLEX (Beckman Coulter, cytoflex B4-R1-Vo) and the data were analyzed using CytExpert software.

### Quantitative polymerase chain reaction (qPCR)

Total RNA was extracted using TRIzol reagent (Invitrogen), and evaluated using a NanoDrop 2000 spectrophotometer (Thermo Scientific, ND-2000). cDNA was synthesized using RevertAid First Strand cDNA Synthesis Kit (Thermo, MA, USA). To assess the microglia signature in NEL-MG, we analyzed the expression of *Tmem119* (Qiagen, Germany, PPM28876A), *P2ry12* (PPM04913C), *Mertk* (PPM34425A), *Tgfb1* (PPM02991B), *Mafb* (PPM05266A), *Gpr34* (PPM04860A), *Pros1* (PPM31106A), *C1qa* (PPM24525E), *Gas6* (PPM05523A), *Csf1r* (PPM03625F), *Hexb* (PPM27125A), and *Gapdh* (PPM02946E) genes. Other primer information (*Trem2, iNOS, TNF-α, IL-1β, IL-6*, and *CCL3*) is indicated in Table S2.

### Immunocytochemistry

Microglial cells were seeded on glass cover slips in 24-well plates. Cells were washed with PBS and cultured; the cultured cells were then fixed in 4% formaldehyde and permeabilized with 0.1% Triton X-100 for 5–10 min. Indirect immunofluorescence was performed using the following primary antibodies: rabbit anti-PU1 (1:200, Abcam, MA, USA; Ab88082), mouse anti-Ki67 (1:500, BD Pharmingen; 550609), mouse anti-F4/10 (1:200, Abcam; Ab6640), rabbit anti-CX3CR1 antibody (1:200, Abcam), rabbit anti-TMEM119 antibody (1:200, Abcam), rabbit anti-IBA-1 antibody (1:500, Wako, MA, USA), and goat anti-IBA1 antibody (1:500, Abcam; Ab48004). Cells were incubated with the primary antibodies diluted in 0.5% Triton X-100 in PBS containing 5% normal donkey serum at 4 °C overnight. After rinsing thrice with PBS for 5 min, Alexa 488- or Alexa-594-conjugated secondary antibodies (Abcam) were used for detection. Nuclei were counterstained with 4′6-diamidino-2-phenylindole (DAPI; Sigma). Cells without the addition of primary antibodies served as negative controls. Fluorescent images were taken using a confocal microscope (LSM 700, Carl Zeiss, Jena, Germany).

### Purification and labeling of synaptosome

One hemisphere (male C57BL/6, Orient Bio Inc. Seoul, Korea), excluding the cerebellum, (∼ 200 – 400 mg), was homogenized in 10 volumes of Syn-PER Synaptic Protein Extraction Reagent (Thermo Fisher Scientific, Part No. 87785) using a 7 mL Dounce tissue grinder with 10 up-and-down even strokes. The homogenate was centrifuged at 1,200 × *g* for 10 minutes to remove the cell debris, and the supernatant was centrifuged at 15,000 × *g* for 20 minutes to obtain the synaptosome pellet. The pellets were gently resuspended in the respective reagent. Synaptosomes were conjugated with an amine-reactive dye (pHrodo Red, SE; Thermo Scientific; #P36600) in 0.1 M sodium carbonate (pH 9.0) at room temperature. After 2 h of incubation, unbound pHrodo was washed-out via multiple rounds of centrifugation and pHrodo-conjugated synaptosomes were resuspended in isotonic buffer containing 5% DMSO for subsequent freezing (Invitrogen, #P36600).

### Phagocytosis assay

To assess the phagocytic activity, NEL-MG at a density of 2 × 10^5^ cells/mL were seeded on a 12-mm coverslip in 24-well cell culture dishes. NEL-MG were treated with 2 μL of red fluorescent latex beads (2 μm, Sigma-Aldrich, St. Louis, MO, USA), HiLyte™ Fluor 488-labeled amyloid β peptide 25-35 (2 μL), or pHrodo-conjugated synaptosomes for 2 h at 37 °C. HiLyte™ Fluor 488-labeled amyloid β peptide 25-35 (Anaspec, AS-633308) was prepared according to the manufacturer’s protocol. Phagocytic activity was then stopped by adding 2 mL of ice-cold PBS. The cells were washed twice with ice-cold PBS, fixed, stained with a microglial marker (IBA-1), and counterstained with DAPI. The cells were analyzed using confocal microscopy (TCS SP5, Leica) and a DeltaVision fluorescence microscopy system (Applied Precision).

### Scratch wound assay

NEL-MG were seeded onto 24-well plates in a 100% confluent monolayer until they were 95% confluent and were wounded by making a perpendicular scratch with a 200 μL pipette tip. The cells were replenished with fresh growth medium and wound closure was documented by photographing the same region at different times (0–6 h). The wound area was calculated as the open wound area/total area.

### Cytokine profiles

The supernatant was analyzed using a Proteome Profiler Mouse Cytokine Array Panel A Kit (R&D Systems; catalog number ARY006) at the baseline and after LPS stimuli according to the manufacturer’s indications. Images were captured using a LAS 4000 (lmageQuant™) and analyzed using ImageJ software program.

### RNA sequence and data analysis

RNA quality was assessed with an Agilent 2100 bioanalyzer using an RNA 6000 Nano Chip (Agilent Technologies, Amstelveen, Netherlands). The library construction was performed using a QuantSeq 3′-mRNA-Seq Library Prep Kit (Lexogen, Inc., Austria) according to the manufacturer’s instructions. High-throughput sequencing was performed as 75 single-end sequences using NextSeq 500 (Illumina, Inc., USA). QuantSeq 3′-mRNA-Seq reads were aligned using Bowtie2 (Langmead and Salzberg, 2012). Differentially expressed genes were determined based on the counts from unique and multiple alignments using coverage in Bedtools (Quinlan AR, 2010). The Read Count data were processed based on the quantile normalization method using Edge R within R (R Development Core Team, 2016) using Bioconductor (Gentleman et al., 2004). Gene classification was based on searches done using DAVID (http://david.abcc.ncifcrf.gov/) and Medline databases (http://www.ncbi.nlm.nih.gov/).

### Neurospheroid culture

The whole brain was dissected from postnatal 1–2-day-old mice (C57BL/6, Orient Bio Inc. Seoul, Korea). Brain tissues were then cut and chopped in HBSS (Gibco) for 3 min. Next, the dissected brain was centrifuged at 300 × *g* for 5 min, after which the pellet was resuspended and washed twice in D-PBS. To detect the capacity for self-renewal, 10^5^ cells were plated onto each well of a 25T-flask in growth-promoting medium: DMEM/F12 containing B27 supplement (×50), minus vitamin A (12587010, ThermoFisher), 50 ng/mL FGF2 (100-18B, PEPROTECH), and 50 ng/mL EGF (AF-100-15, PEPROTECH). Cultures were maintained at 37 °C in a 5% CO2 incubator for neurospheroid (NS) formation.

### CellTracker labeling

NEL-MG were labeled using CellTrackerRed CMTPX (Invitrogen) before mixing with NS. Labelling was performed according to the manufacturer’s indications.

### Mixing of NEL-MG with neurospheroids

NEL-MG were co-mixed with dissociated NS or post-treated with the already formed NS. NS were mixed with NEL-MG in a 9:1 ratio (1.8 × 10^6^ NS: 2 × 10^5^ NEL-MG) or a 7:3 ratio (1.4 × 10^6^ NS: 6 × 10^5^ NEL-MG) in DMEM/F12 containing B27 supplement (×50), 50 ng/mL FGF2, and 50 ng/mL EGF. After the addition of NEL-MG, the plates were maintained under static conditions in a shaking incubator (70 rpm) at 37 °C with 5% CO2.

### Statistical analysis

The statistical significance of differences between groups was assessed using an unpaired t-test or one-way analysis of variance using GraphPad Prism version 7 for Mac (GraphPad, La Jolla, CA). A *post-hoc* test was performed using one-way analysis of variance when the p-values were significant (p < 0.05).

## Declarations

### Ethics approval and consent to participate

All experimental procedures were approved by the Institutional Animal Care and Use Committee (IACUC) of the CHA University (IACUC200116)

### Consent for publication

Not applicable

### Availability of data and material

All data generated and/or analyzed during the current study are available from the corresponding author on reasonable request.

### Competing interests

The authors declare that they have no competing interests

### Funding

- This research was supported by the National Research Foundation of Korea (NRF) grant funded by the Korea government (MIST) (2019M3C7A1032561), by Basic Science Research Program (2020R1F1A1074668), by a grant of the Korea Health Technology R&D Project through the Korea Health Industry Development Institute (KHIDI), funded by the Ministry of Health &Welfare, Republic of Korea (HI16C1559).

### Authors’ contributions

- You MJ conducted *in vitro* experiments and wrote the manuscript. Rim C analyzed transcriptome. Kang YJ interpreted all data and revised the manuscript. Kwon MS supervised all process, supported experimental conception and design, and approved final submission of manuscript. All authors critically revised the manuscript and confirmed author contribution statement.

## Acknowledgements

Not applicable

## Supplementary figures

**Figure S1.**
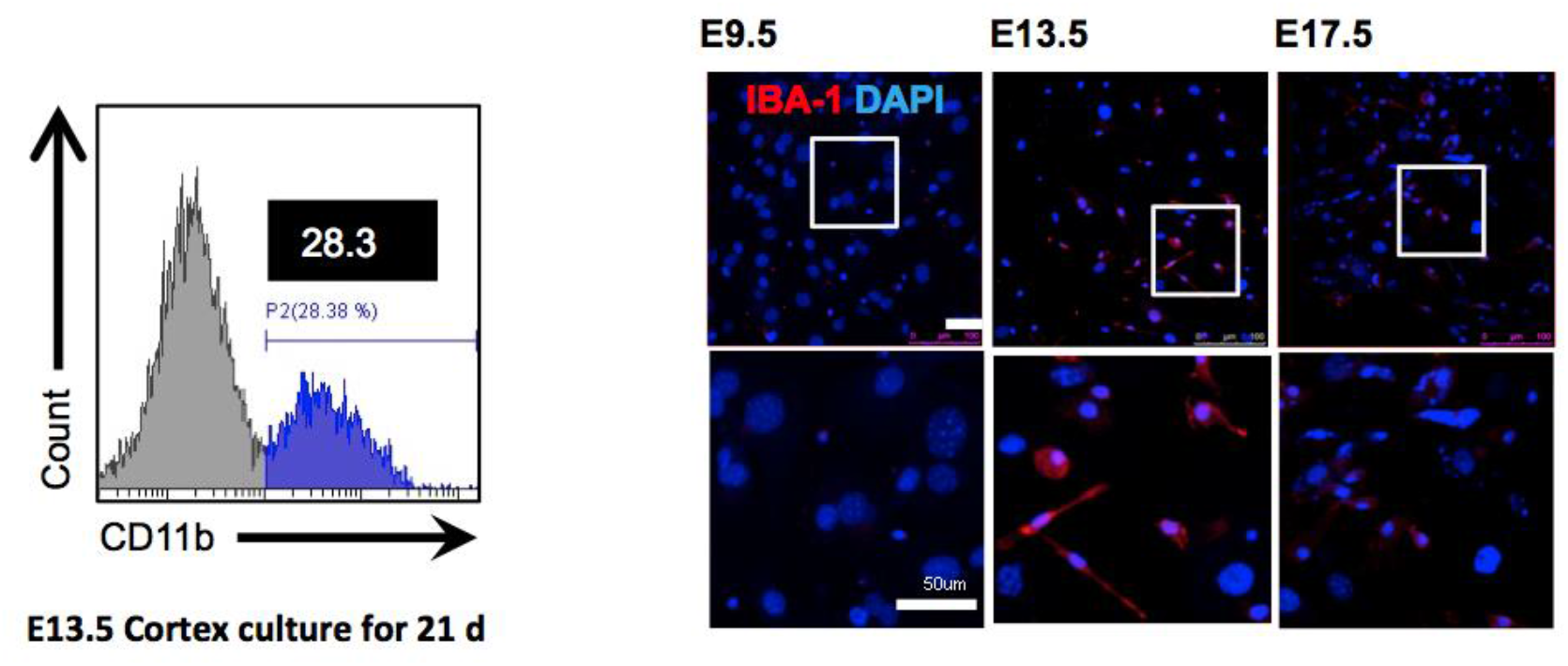
Comparison of the cell yields according to the region and embryo stage. Flow cytometry revealed that the dissected head neuroepithelial layer (NEL) from mouse E13.5 could yield a higher ratio of CD11b-positive cells than the brain cortex separated from an identical mice group. Immunofluorescence showed that 13.5 NEL have a higher number of IBA-1-positive cells than neuroepithelial layer from mouse E9.5 or E17.5, when cultured for 21 days. Scale bar = 50 μm.

**Figure S2.**
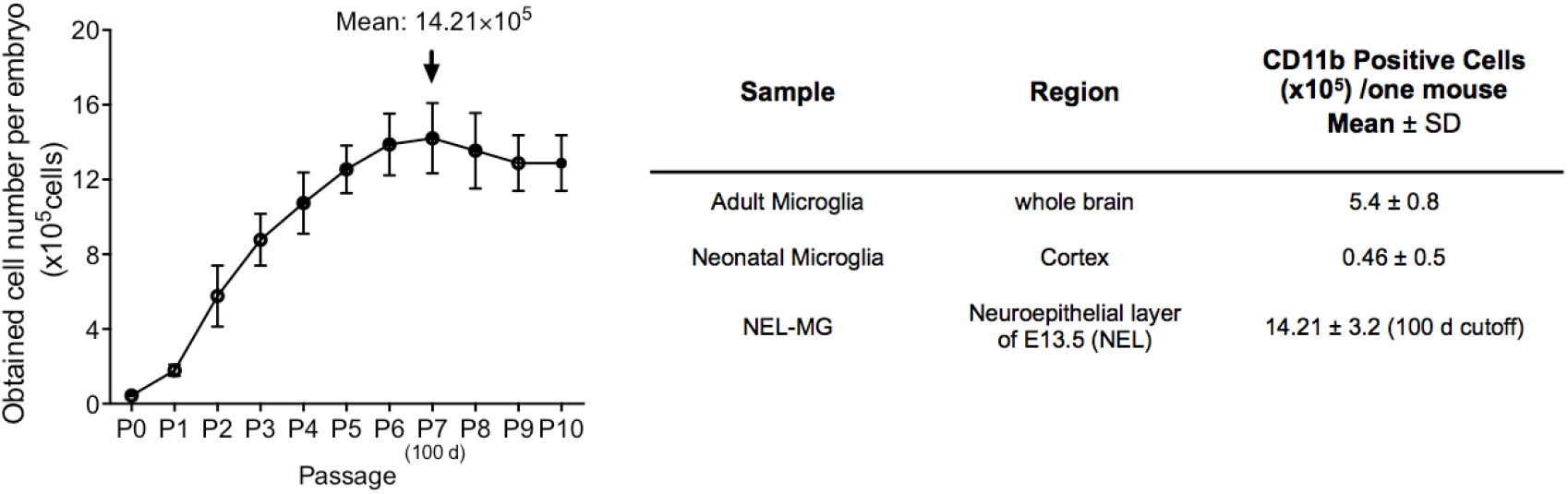
Mass production of NEL-MG via subculture. Our method produced thirty times the number of microglial cells than that of neonatal microglia when we used a cutoff of 100 days. The data are shown as mean ± standard error of the mean (SEM).

**Figure S3.**
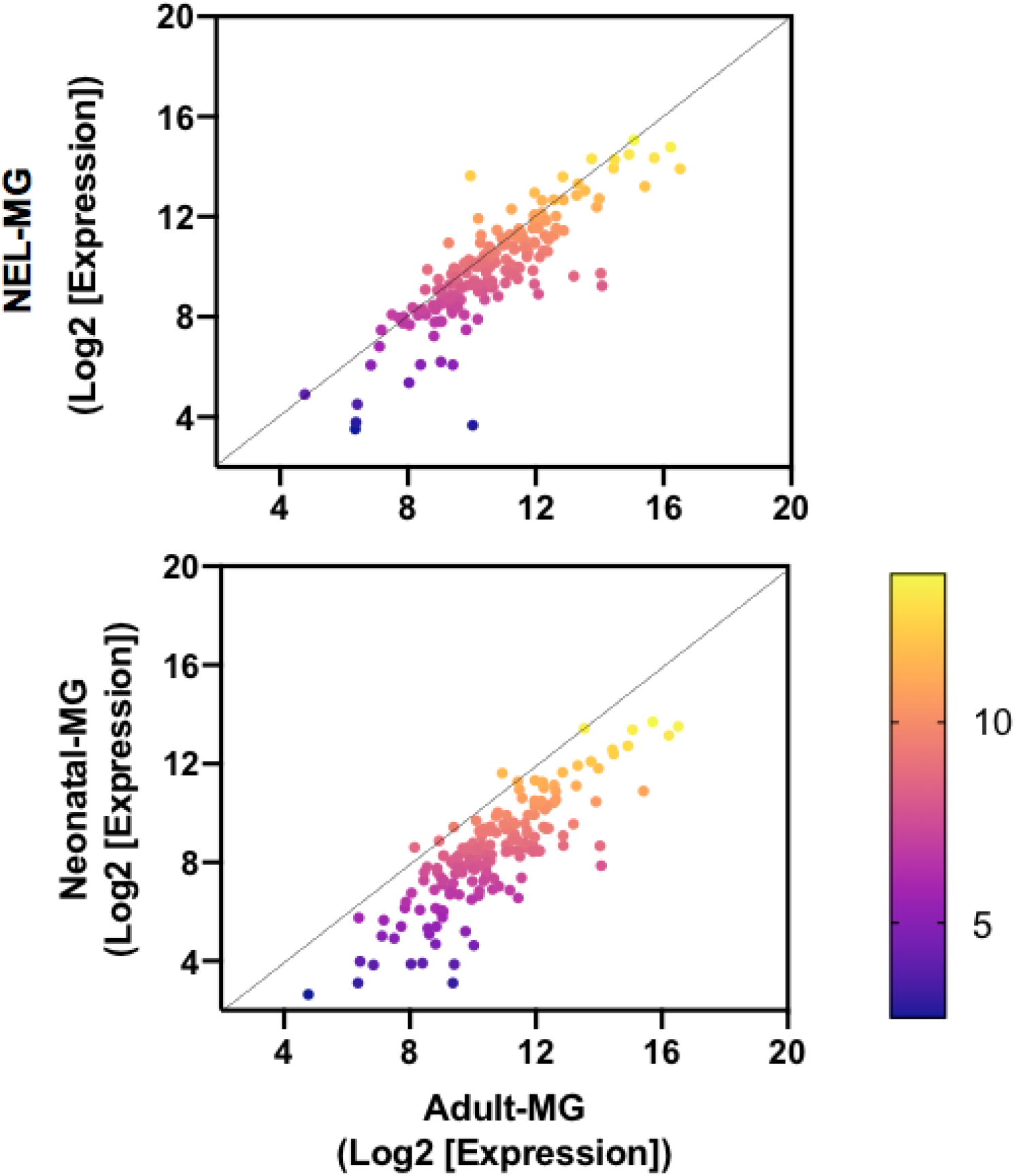
Scatter plot comparison between adult microglia and NEL-MG or neonatal microglia (MG) Diagonal line indicates no significant difference between the two groups (fold change = 1) and the intensity values are normalized to the log2 transformed expression value.

**Figure S4.**
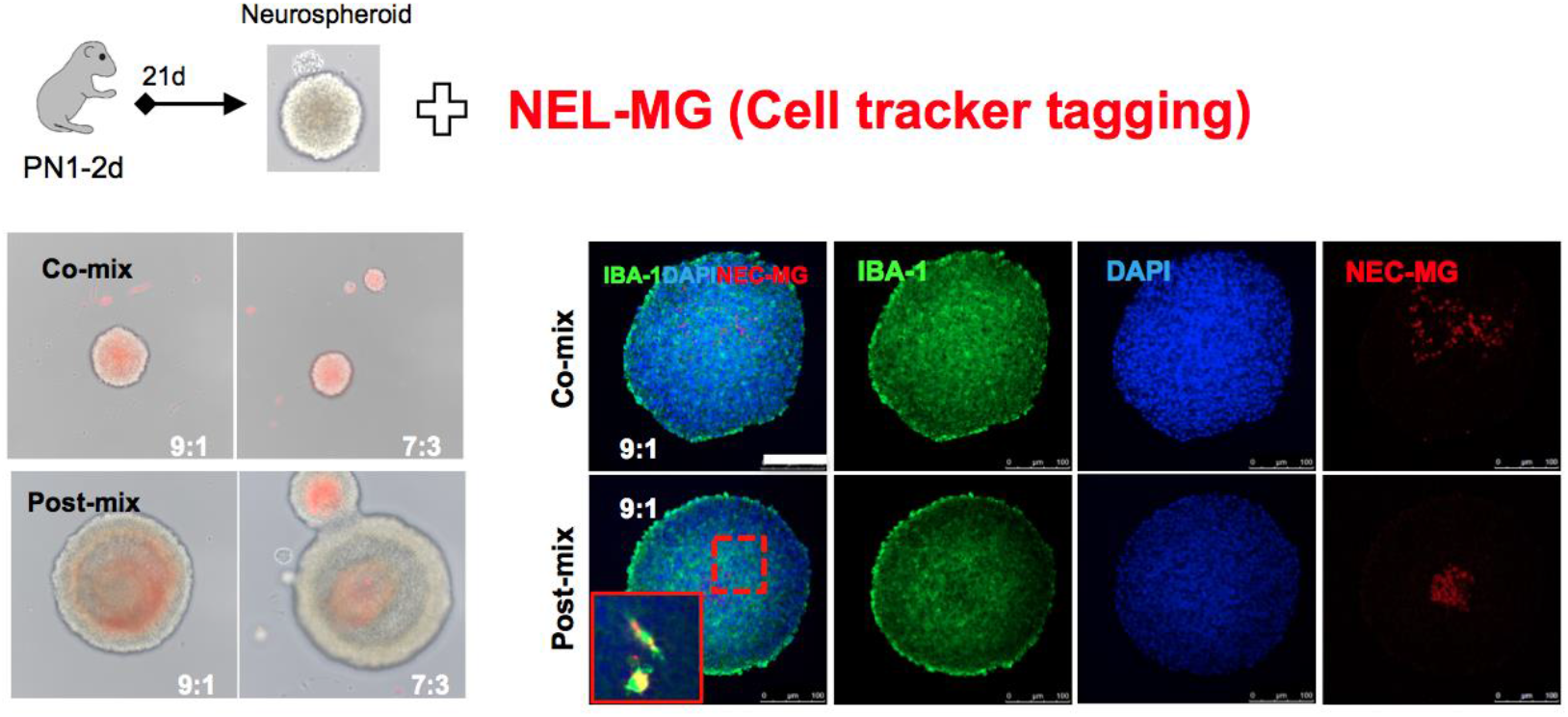
Mix with neurospheroid. CellTracker-tagging NEL-MG (red) were mixed with neurospheroids (NS) at different ratios and times. NEL-MG were mixed evenly with NS when we co-mixed them. IBA-1 can label both resident microglia and mixed NEL-MG.

**Table S1.**
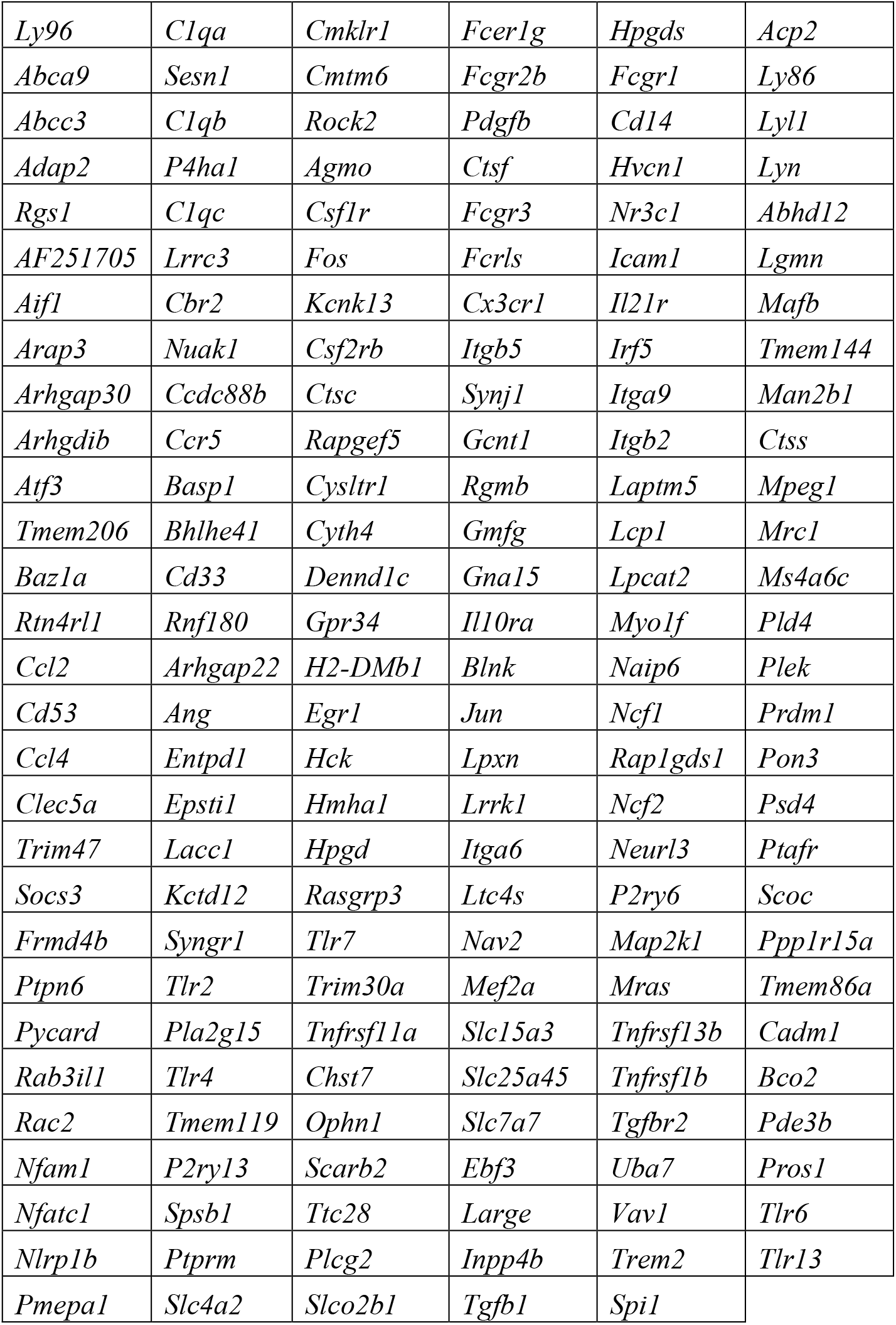
List of microglial signature genes

**Table S2.**
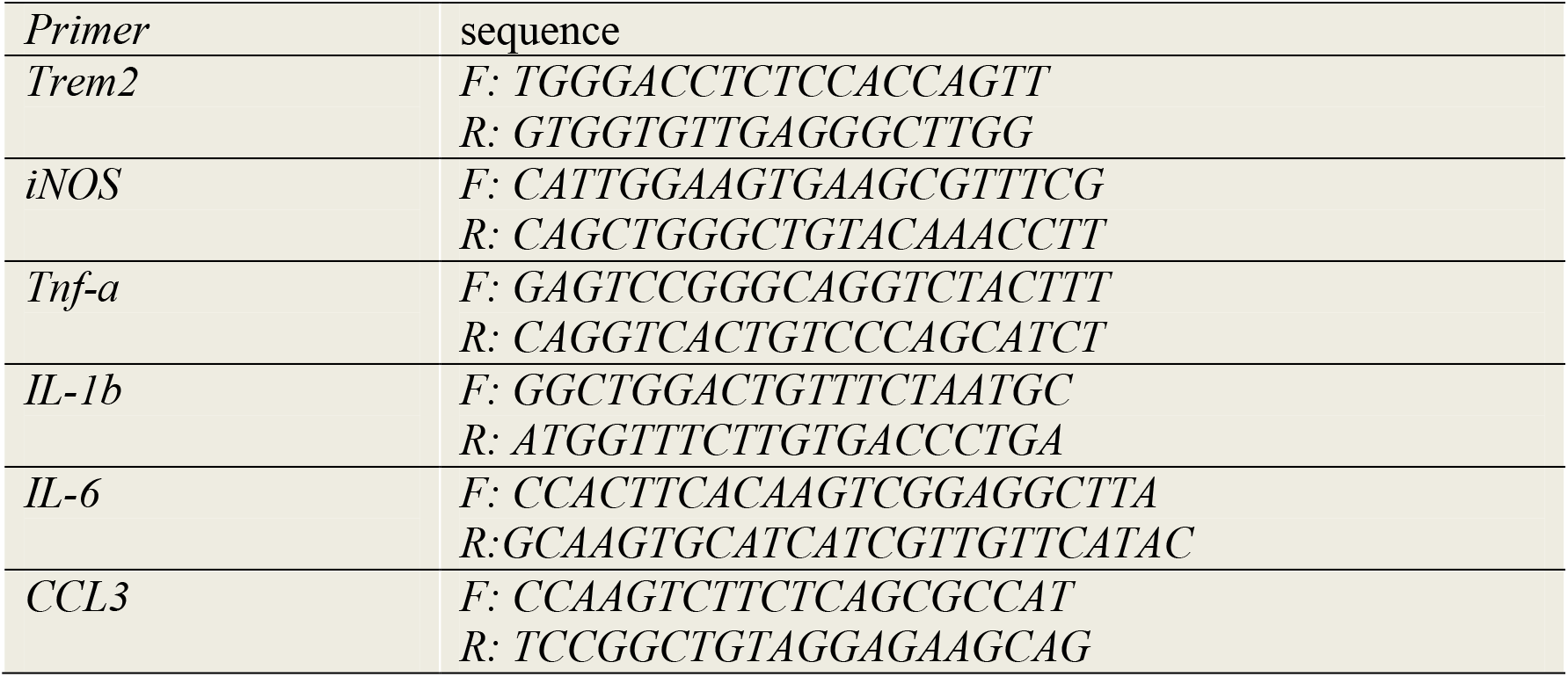
Primers information.

## Notes

### Competing Interest Statement

The authors have declared no competing interest.

